# Interpretive Summary. Heat detection alerts and LH surge. Adriaens

**DOI:** 10.1101/249169

**Authors:** Ines Adriaens, Wouter Saeys, Chris Lamberigts, Mario Berth, Katleen Geerinckx, Jo Leroy, Bart De Ketelaere, Ben Aernouts

## Abstract

Both estrus detection and timely insemination are important factors in optimizing fertility management. The latter is dependent on ovulation time, which is preceded by the LH surge. The performance of an estrus detection system based on activity and based on milk progesterone was evaluated and the timing of the alerts was contrasted against the moment of the LH surge. Activity alerts had a sensitivity of 83% and a positive predictive value of 66%; the LH surge followed average 9.4 ± 16.1 hours later. Using milk progesterone, one can reliably detect luteolysis, which is followed by the LH surge after 62 ± 12 hours.

---

Both maximal detection of estrus and timely insemination are important to improve on-farm fertility performance (Hockey et al., 2010a). Although the latter is strictly associated with ovulation time (**OT**), the link between OT and estrus alerts provided by automated estrus detection systems is seldom researched due to the high workload and costs of these studies (Roelofs, 2005). In this study, the sensitivity and positive predictive value (**PPV**) of an activity- and a progesterone (**P4**)-based system (period 1+2), and subsequently, their link with the LH surge preceding ovulation were investigated (period 2).

The study was approved by the ethical committee of KU Leuven (ID P010/2017) and conducted in an experimental dairy farm in Flanders, Belgium. Twenty-two regularly cycling Holstein-Friesian cows, aged 3.4±1.2 years (mean±SD) with parities 1 to 4 and being 30 to 293 days in milk (**DIM**) at the start of the study were included. The cows were milked with an automated milking system (VMS, Delaval, Tumba, Sweden) and were fed with a mixed ration of grass and corn silage and thereby supplemented with concentrates. During 53 days, a mixed milk sample of each milking was taken automatically by the sampling unit (VMX, Delaval, Tumba, Sweden), following the procedure of the dairy herd improvement protocol prescribed by ICAR (ICAR International committee for animal recording, 2014) and stored at -20°C. At the end of the trial, the certified lab MCC-Vlaanderen (Lier, Belgium) determined the P4-concentration using a Ridgeway ELISA kit, for which more details are described in Adriaens et al., (2017). The P4 concentration of each sample was also measured automatically on-farm by the Herd Navigator^TM^ system (Lattec, Delaval, Hiller□d, Denmark). This device allowed to estimate the moment at which the cows possibly came in estrus without the need for synchronizing the ovaries. Based hereon, a reference estrus period (**REP**) was defined as a period of low milk P4 (< 5 ng/mL) of minimum 5 and maximum 10 days, in which 15 to 50 hours after the start of the REP a preovulatory follicle of at least 13 mm was detected by an expert veterinarian using a transrectal ultrasound scanner (A6v, Sonoscape Medical Corp., Shenzhen, China) (Hockey et al., 2010b; a). This methodology allowed us to identify all estrous periods of the cows, including silent estruses, while avoiding the need of daily ultrasound examinations. The sensitivity of each estrus detection method was defined as the number of times an alert was given by a system during the REP (true positive, **TP**) divided by the total number of REP (true estrus). The PPV was the number of TP compared to the total number of alerts for that method.

Each cow was fitted with a commercial activity meter (ActoFIT, version 2015, Delaval, Tumba, Sweden) on the neckband. Increased activity was monitored via the algorithm included in the DelPro Farm Manager software. This algorithm generated activity alerts in 3 possible levels (+/++/+++) dependent on the actual restlessness of the cow compared to her normal behavior. In this study, only the highest level of activity within the REP was considered. Each alert not associated with a REP was considered a false alarm (**FP**). Progesterone-based alerts were generated with the recently developed on-line monitoring system named *‘*P4 Monitoring Algorithm using Synergistic Control’ (**PMASC**), consisting of a mathematical model describing the different parts of the P4 profile (Adriaens et al., 2017) and a statistical process control chart to indicate luteolysis (Adriaens et al., 2018). An alert was taken as TP if followed immediately by a REP within 24 hours. All other attentions were considered as FP. The sensitivity and PPV of the different estrus detection systems were determined for a period of 41 days (i.e. period 1+2), allowing for a training period of 12 days for PMASC in which only P4 samples were taken.

The preovulatory LH surge was monitored as a proxy for ovulation (period 2) between day 28 and 53 of the trial. If the presence of a preovulatory follicle was confirmed after a P4 drop, two-hourly blood samples were collected from the jugular vein over a period of 72 hours, starting 36 hours after the moment the raw milk P4 started a consistent drop towards concentrations under 5 ng/mL. A preliminary study showed that this amount of samples was needed to ensure a full image of all LH surges, but if the cow showed post-estrous bleeding, sampling was stopped. The clotted blood samples were centrifuged at 2300 *g* to collect the serum, from which 3 aliquots were stored at -20°C. Seven to 11 days later, the disappearance of the preovulatory follicle and the presence of a corpus luteum (**CL**) were verified by ultrasonography. At the end of the trial, the serum LH concentration was measured on a BEP2000 system (Siemens Healthcare Diagnostics, Marburg, Germany) with a commercially available bovine LH ELISA kit (Abnova, Taipei City, Taiwan), having a within-run coefficient of variation of 6.3% (average LH 13.2 ng/mL) and 5.5% (average LH 45.2 ng/mL) and a limit of quantification of 3 ng/mL. The presence of an LH surge was visually determined on a time versus LH concentration graph, and the moment of the LH surge taken as the maximal LH concentration, which was between 17 ng/mL and 49 ng/mL. In all cases, this surge concentration was more than 8-fold the baseline LH concentration. In total, LH samples were taken during 24 REP from 22 cows in 25 days. However, to obtain unbiased results, cows with severe health or fertility problems known to affect endocrinology (Dobson et al., 2008; Walker et al., 2008), were excluded for this part of the study. These included one animal treated for milk fever, one being severely lame and one with endometritis, and also three cows treated for a luteal cyst and three animals with a follicular cyst or that went in anestrus after a normal cycle. A total of 15 REP (9 primiparous, 6 multiparous cows, 158 ± 53 days in lactation, body condition score 3.2 ± 0.3, mean ± SD) remained for the analysis.

The time interval (**TI**) between each true positive activity or P4 attention and the moment of maximal LH concentration was calculated in hours, further referred to as **TI_ACT_** and **TI_PMASC_** respectively. Additionally, it was evaluated how the LH surge related with a number of model-based indicators derived from PMASC. Hereto, a TI was calculated with (1) the inflection point of the decreasing Gompertz function describing the luteolysis (**TI_IP_**); (2) the intercept of the tangent line at the inflection point with the time-axis (**TI_IC_**); (3) the moment that the model surpassed a fixed threshold of 3, 5, 7 or 10 ng/mL (**TI_TMOD_X_**); and (4) the moment the model undercut 85, 90 or 95% of the maximum P4 model concentration minus the baseline (**TI_TB_X_**), with X representing the respective percentages and thresholds (Adriaens et al., 2017, 2018).

Thirty-five REP were detected in 22 cows during period 1+2. The number of alerts given per system, together with their sensitivity and PPV, are summarized in Table 1. The activity alerts are expressed according to their maximum activity level of respectively ‘+’, ‘++’ or ‘+++’ for each REP. When all three activity levels were considered, a sensitivity of nearly 83% was obtained. However, estrus detection solely based on the ‘+’ activity level is not reliable, as 13 of the 19 ‘+’ attentions (68.4%) were false alarms (PPV of 31.6%). These moderate increases in activity are most likely to be caused by dominance fighting or environment related factors. Using only the ‘++’ and ‘+++’ attentions decreased the number of false alarms to 8.0% and 0.0% respectively, but also reduced the sensitivity to 65.7% and 40.0%. The main advantage of activity meters over traditional visual estrus detection is the automated and continuous nature of the system which limits the time required for estrus detection by the farmer. However, the sensitivity and PPV in this study did not meet the standard required for sensor systems, and moreover, the current results only include the cows verified to be cycling and healthy at the start of the trial. Accordingly, the false alarm rate for ‘++’ and ‘+++’ attentions might further increase when including e.g. nymphomaniac animals with follicular cysts. Additionally, activity scoring does not identify true silent ovulations (17.1% in our study).

**Table 1.** Summary of the attentions, sensitivity, positive predictive value and time intervals (TI_X_) for the different estrus detection systems

| REP<sup>1</sup> | Period 1+2<br>35 |  |  |  |  | Period 2<br>15 |  |  |
| --- | --- | --- | --- | --- | --- | --- | --- | --- |
|  | P<sup>5</sup> | TP<sup>6</sup> | FP<sup>7</sup> | Sensitivity | PPV<sup>8</sup> | TP<sup>6</sup> | Mean ± SD | TI<sub>X</sub><sup>3</sup><br>Range [min ; max] |
| Activity attentions | 44 | 29 | 15 | 82.9% | 65.9% | 12 | 9.4 ± 16.1 | 47.0 [-8.0 ; 39.0] |
| + | 19 | 6 | 13 | 82.9%<sup>4</sup> | 31.6% | 2 |  |  |
| ++ | 11 | 9 | 2 | 65.7%<sup>4</sup> | 81.8% | 5 |  |  |
| +++ | 14 | 14 | 0 | 40.0%<sup>4</sup> | 100.0% | 5 |  |  |
| PMASC<sup>1</sup> | 35 | 35 | 0 | 100.0% | 100.0% | 15 | 62.0 ± 12.2 | 35.7 [45.9 ; 81.6] |
| IP<sup>8</sup> |  |  |  |  |  |  | 73.4 ± 11.1 | 35.2 [52.0 ; 87.2] |
| IC<sup>9</sup> |  |  |  |  |  |  | 55.7 ± 12.4 | 40.6 [38.7 ; 79.3] |
| TB85<sup>10</sup> |  |  |  |  |  |  | 56.4 ± 11.2 | 34.6 [41.8 ; 78.4] |
| TB90<sup>10</sup> |  |  |  |  |  |  | 46.2 ± 15.5 | 43.9 [21.6 ; 75.5] |
| TB95<sup>10</sup> |  |  |  |  |  |  | 43.0 ± 17.2 | 66.4 [4.9 ; 71.3] |
| TMOD<sub>10</sub><sup>11</sup> |  |  |  |  |  |  | 66.6 ± 11.1 | 32.9 [48.3 ; 81.2] |
| TMOD<sub>7</sub><sup>11</sup> |  |  |  |  |  |  | 61.6 ± 11.4 | 34.7 [45.6 ; 80.3] |
| TMOD<sub>5</sub><sup>11</sup> |  |  |  |  |  |  | 56.8 ± 12.6 | 44.2 [35.1 ; 79.3] |
| TMOD<sub>3</sub><sup>11</sup> |  |  |  |  |  |  | 47.8 ± 18.1 | 63.5 [13.9 ; 77.4] |

<sup>1</sup> REP: Reference estrus period; <sup>2</sup>PMASC – Progesterone (P4)-based monitoring algorithm using Synergistic Control; <sup>3</sup> TI<sub>X</sub>: time interval with alert or model-based indicator and the preovulatory LH surge; <sup>4</sup> The sensitivity for the activity alerts was calculated cumulatively, using all (+/++/+++)<sup>4</sup> true positives for '+', only the '++' and '+++' for '++' and only the '+++' alerts for '+++'; <sup>5</sup> Positives, estrus alerts generated with PMASC or the activity based system; <sup>6</sup> True positives: alerts associated with a REP; <sup>7</sup> False positives: alerts not associated with a REP; <sup>8</sup> IP = inflection point, <sup>9</sup> IC = intercept tangent line at IP and t-axis; <sup>10</sup> TB\_X: threshold based on model passing X% of the maximal + baseline P4 value; <sup>11</sup> TMOD\_X: model passing a threshold of X ng/mL;

The PMASC system identified all 35 REP in the 41 day study period and did not given any false alerts (sensitivity and PPV of 100%). In a large study conducted by Friggens et al., (2008), comparable sensitivities of 93.3% to 99.2% were reached, using the model described by Friggens and Chagunda, (2005). This result might be influenced by the way the REP was defined. However, the additional check-up of the ovaries using ultrasonography aided to confirm each estrous period, and physiology dictates that estrus cannot occur during luteal phases of the cycle. Therefore, the current method applied is a watertight one, and no better alternative is available today.

Typically, health affects the extent of restlessness shown during estrus, which influences the performance of the activity-based systems. However, as long as there is no better practice-based insight in when it is recommended to inseminate these ‘unhealthy’ cows, or which factors (such as body condition score, diseases, insufficient uterus tonus, …) should be taken into account, the value of P4-based systems might be over-estimated considering the above-mentioned performance. To avoid that health problems could affect the results of the second part of this study in which the link between the LH surge and the respective alerts given was investigated, only REP known to originate from healthy cows were included (period 2, Table 1).

The baseline concentration of serum LH and the LH surge was respectively 1.46±0.84 ng/mL and 36.3±11.3 ng/mL (both mean±SD) and the latter developed and faded for all cows within 8 hours (= 4 measurements). An overview of the LH data and the P4 profiles centered around the LH surge is shown in Figure 1. The TI between this LH surge and the different alerts (TI_ACT_, TI_PMASC_, TI_IC_, TI_IP_, TI_MOD_X_, TI_TB_X_) is given in Table 1. Three REP could not be associated with any activity attention and from the other 12, 2 had only a ‘+’ alert, which was shown before to be unreliable for estrus detection. The range from alert to LH surge varied from 39 hours before the LH surge to 8 hours after it, and had a SD of 16.1 hours. Roelofs and colleagues reported that ovulation occurred 29.3±3.9 hours after the onset of an increased number of steps (TI between 22-39 hours) and 19.4±4.4 hours after the end of the increased number of steps (TI between 12-35 hours) (Roelofs et al., 2005), which shows that alternative sensor systems and data processing algorithms might improve the results. However, evaluating these was outside the scope of this study.

**Figure 1.**
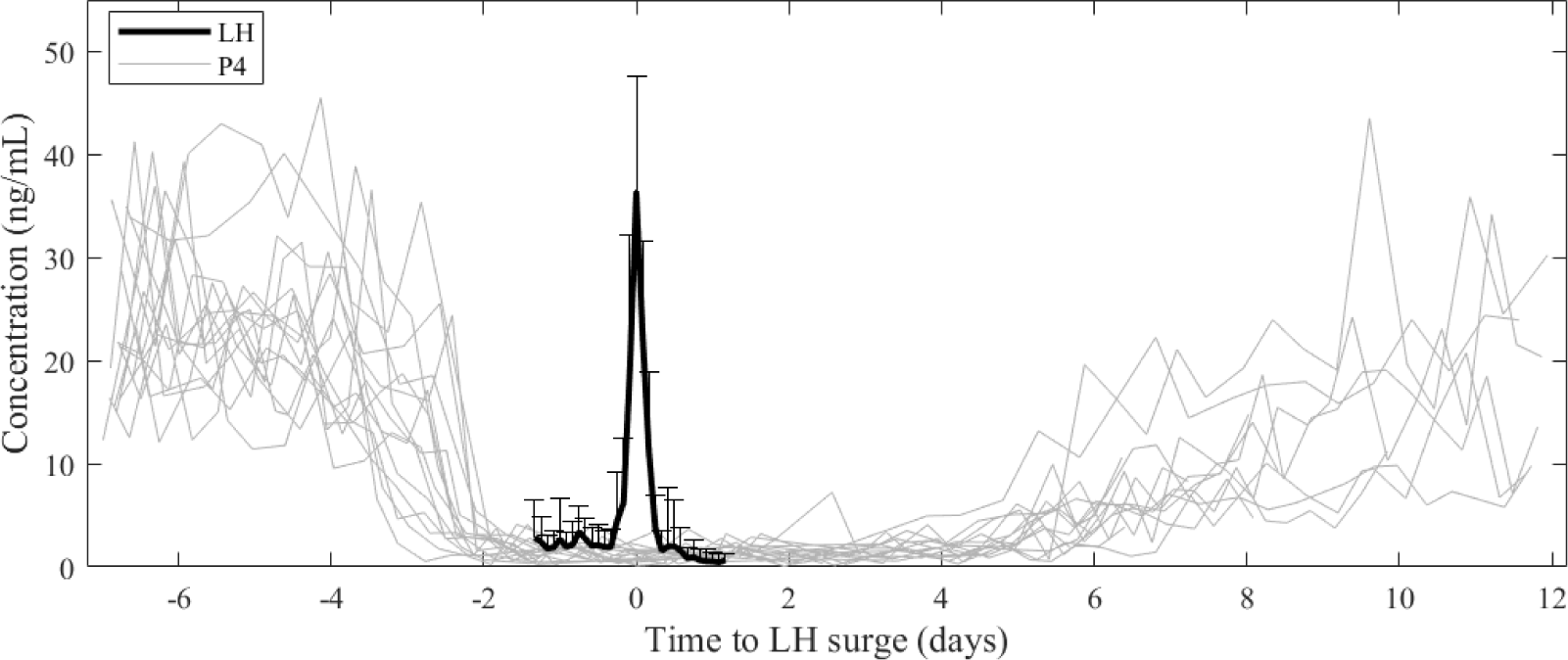
Overview of the progesterone (P4) profiles centered around the LH surge. A large variability in the P4 profiles is noted both before and after the LH surge, but all cows had an active corpus luteum, reflected in a P4 concentration higher than 5 ng/mL within 7 days after the LH surge.

The average TI from luteolysis detected with the PMASC system to the LH surge was 62 hours, with a minimum of 46 and a maximum of 82 hours, resulting in a range of 36 hours and a SD of 12.2 hours. Both IP, TB85, Tmod_10_ and Tmod_7_ perform similarly (range TI_X_ of 33 – 35 hours, SD 11.1 to 11.4 hours). With a luteal P4 concentration between 20 and 30 ng/mL and a follicular concentration of approximately 2.5 ng/mL, TB85 represents the moment that the model goes below 2.5 + 0.15 * (20 - 2.5) = 5.1 to 2.5 + 0.15 * (30 - 2.5) = 6.6 ng/mL. Using the TB85 indicator, and thereby taking into account the drop compared to the absolute maximal P4 concentration, resulted in a lower TI range than when a fixed threshold of 5 ng/mL was used on the model (TI_TMOD5_). However, when a fixed threshold of 10 on the model was used (TI_TMOD10_), an even smaller range of 33 hours was noted (SD 11.1 hours). These results demonstrate that the start of luteolysis might be more indicative for the LH surge than the P4 being fully cleared from the cow’s bloodstream. In the current study, samples were taken each milking. However, when less samples are taken, it is expected that the model-based indicators will outperform the luteolysis alerts of PMASC. Using model-dependent rather than data-dependent guidelines might therefore make the monitoring system more robust. Furthermore, they can be calculated for each profile, allowing the flexibility to account for differences in shapes and absolute levels, and thereby providing the possibility to further optimize insemination advice, e.g. by fixing the slope-determining parameters to physiologically relevant ranges when less frequent samples are available (Meier et al., 2009).

The limited number of cases and the fact that only one farm was included in the study bring along that no steady claims can be made on the optimal insemination time for attentions generated with these systems, nor can the observations and results be extended to genetically different herds. Therefore, more data should be collected to confirm or contradict our results, taking into account extended measures of physiology, genetics, management, etc. Unfortunately, studies taking ovulation time into account are time consuming, labor intensive and expensive. The current smaller type of study allows to place larger studies, which typically only evaluate one estrus detection system in absence of a good reference for ovulation into perspective. We therefore identify the need for additional research including both different estrus detection tools and a good reference for the moment of ovulation. For the latter, monitoring LH in the blood serum allows to spread the workload as the analysis and interpretation of the samples can be organized post-hoc. This circumvents the need for applying a synchronization protocol, so the fertility of the animals can be assessed in a natural, farm-representative setting. Moreover, the blood sampling is considered to cause less stress and have no direct impact on the ovaries, compared to ultrasound rectal scanning of the cows. Additionally, to improve precision, ultrasound scanning and interpretation of the images should preferably be done by the same persons for the full study. This is, however, not feasible when cows are not synchronized.

To conclude, we found that P4 monitoring allows for accurate identification of luteolysis which outperforms the activity-based estrus detection system in terms of sensitivity and PPV. Moreover, using LH as a proxy for ovulation, the P4-based system and derived model-indicators seem to allow for a more reliable identification of the correct moment for insemination.

## ACKNOWLEDGEMENTS

This work was supported by the Institute for the Promotion of Innovation through Science and Technology in Flanders, Belgium (IWT) [IWT-LA project 110770]. Ines Adriaens and Ben Aernouts are supported by the Fund for Scientific Research (FWO) Flanders, respectively grant number 11ZG916N and 12K3916N. We thank Shahbaz Bashir and Mathilde Tinel for their assistance during the blood sampling. Lattec (Hillerød, Denmark) supported this study by supplying additional P4 sticks and modifying the Herd Navigator software to take samples at every milking event.

